# Increased transmission of SARS-CoV-2 lineage B.1.1.7 (VOC 2020212/01) is not accounted for by a replicative advantage in primary airway cells or antibody escape

**DOI:** 10.1101/2021.02.24.432576

**Authors:** Jonathan C. Brown, Daniel H. Goldhill, Jie Zhou, Thomas P. Peacock, Rebecca Frise, Niluka Goonawardane, Laury Baillon, Ruthiran Kugathasan, Andreia L. Pinto, Paul F. McKay, Jack Hassard, Maya Moshe, Aran Singanayagam, Thomas Burgoyne, the ATACCC Investigators, PHE Virology Consortium, Wendy S. Barclay

## Abstract

Lineage B.1.1.7 (Variant of Concern 202012/01) is a new SARS-CoV-2 variant which was first sequenced in the UK in September 2020 before becoming the majority strain in the UK and spreading worldwide. The rapid spread of the B.1.1.7 variant results from increased transmissibility but the virological characteristics which underpin this advantage over other circulating strains remain unknown. Here, we demonstrate that there is no difference in viral replication between B.1.1.7 and other contemporaneous SARS-CoV-2 strains in primary human airway epithelial (HAE) cells. However, B.1.1.7 replication is disadvantaged in Vero cells potentially due to increased furin-mediated cleavage of its spike protein as a result of a P681H mutation directly adjacent to the S1/S2 cleavage site. In addition, we show that B.1.1.7 does not escape neutralisation by convalescent or post-vaccination sera. Thus, increased transmission of B.1.1.7 is not caused by increased replication, as measured on HAE cells, or escape from serological immunity.

## Introduction

In late 2019, SARS-CoV-2 emerged into humans from animals and rapidly led to a global pandemic. In September 2020, a new variant of SARS-CoV-2, lineage B.1.1.7 (Variant of Concern 202012/01) emerged in the UK (Rambaut et al., 2020). B.1.1.7 is distinguished by a large number of mutations and a long phylogenetic branch length separating it from its closest sequenced isolates. The genetic distance from other viruses has prompted suggestions that B.1.1.7 may have evolved during extended infection of an immunocompromised host (Rambaut et al., 2020). In late 2020 and early 2021, B.1.1.7 spread rapidly to become the dominant lineage in the UK. This is likely accounted for by increased transmissibility measured by an increase in the effective reproduction number (R_t_) of 0.4-0.7 (Volz, Mishra, et al., 2021). B.1.1.7 has now been detected in 88 other countries, and has become predominant in several of these (O’Toole et al., 2021). Moreover, recent reports suggest that B.1.1.7 infection results in an approximately 70% higher hazard of death compared to other strains (Challen et al., 2021; Davies et al., 2021). To effectively control SARS-CoV-2 and to assess the risk of future variants, it is vital to understand the phenotypic characteristics and the underpinning mutations which have resulted in the higher transmissibility and pathogenicity of the B.1.1.7 lineage.

The B.1.1.7 lineage is characterised by 23 mutations across the viral genome (Rambaut et al., 2020). The spike glycoprotein (S) harbours 9 of these including N501Y, Δ69-70, Δ144 and P681H. N501Y lies in the receptor binding domain (RBD) and has been shown to enhance binding of S to its receptor ACE2 (Starr et al., 2020; Supasa et al., 2021). This mutation gives the B.1.1.7 UK variant the alternative designation 20I/501Y.V1 and has also been observed in several other lineages including the B.1.351 (501Y.V2) South African variant and the Brazilian P.1 (20J/501Y.V3) variant (Faria et al., 2021; Tegally et al., 2020). Two deletions, Δ69-70 and Δ144, map to the N-terminal domain (NTD). Deletions around position 144 have been observed during extended replication *in vitro* and within-host evolution in immunocompromised patients, and have also been linked to escape from NTD-targeting antibodies (Andreano et al., 2020; Choi et al., 2020; Kemp et al., 2021; McCallum et al., 2021; McCarthy et al., 2020). Δ69-70 has arisen in multiple lineages and its effect is unknown but may compensate for a putative fitness cost of substitutions in the RBD (Kemp et al., 2020). P681H is at the S1/S2 cleavage site and could affect the efficiency of furin cleavage. We and others have previously shown that efficient cleavage of S at this site enhances transmissibility and pathogenicity of SARS-CoV-2 (Johnson et al., 2021; Peacock et al., 2020; Zhu et al., 2021). B.1.1.7 also harbours mutations of interest in other genes including a premature stop codon in ORF8, an accessory gene that likely enables immune evasion (Zhang et al., 2020), and a 3 amino acid deletion in NSP6, one of several proteins associated with virus regulation of the innate immune response (Xia et al., 2020). Interestingly, truncation of ORF8 and the identical deletion in NSP6 are also present in several other emerging SARS-CoV-2 variants suggesting a profound degree of convergent evolution even outside the S gene, however their phenotypic effects remain undefined.

The appearance and rapid spread of the B.1.1.7 variant beyond the UK is clearly not a result of chance, for example a founder effect, but due to a transmission advantage conferred by its particular genetic constellation. However, transmission is a multifactorial phenotype and it is not yet clear which of the B.1.1.7 mutations, or specific combination of mutations, contribute to the modification of viral traits which support its increased transmissibility. Improved transmissibility may owe to a combination of factors including more rapid viral replication within a host increasing the amount or duration of virus emitted from an infected host, increased environmental stability of infectious virus, more efficient entry into host cells, improved innate immune evasion that would increase the chance of an exposure leading to infection, or the ability to overcome convalescent and post-vaccination sera thereby increasing the size of the population susceptible to infection.

Epidemiological data to support some of these potential explanations for increased transmissibility are thus far mixed. The results of analyses of Ct values or number of mapped sequencing reads as a proxy for viral load are currently conflicting making it difficult to conclude whether there is a replicative advantage for B.1.1.7 in-host (Golubchik et al., 2021; Kidd et al., 2020; Walker et al., 2021). Another recent study showed similar peak viral burden but an increased duration of infection with B.1.1.7 infection, albeit with a small sample size (Kissler et al., 2021).

The question of whether the B.1.1.7 variant escapes pre-existing immunity is also unclear. In vitro passage of SARS-CoV-2 in the presence of neutralising antibodies can give rise to spike mutations which evade antibody immunity (Andreano et al., 2020; Weisblum et al., 2020) making surveillance for such escape mutations in circulating viruses and measurement of their effects *in vitro* a priority. In an intense burst of research around this question, convalescent and post-vaccination sera have been used in neutralisation assays with B.1.1.7 live virus isolated from infected patients, recombinant viruses generated by reverse genetics and virus pseudotypes (PV) with some or all of the B.1.1.7 mutations in S, but results have been inconsistent. In some studies, PV bearing B.1.1.7 and wildtype S showed equivalent (<2-fold) neutralisation by convalescent sera (Collier et al., 2021; Rees-Spear C et al., 2021) or sera raised against vaccines (Muik et al., 2021; Wu et al., 2021). However, other studies found a modest decreased susceptibility (up to 4-fold) of B.1.1.7 PV to convalescent sera (Hu et al., 2021; Wang et al., 2021) or vaccine sera (Collier et al., 2021). Sera raised against BNT162b2 vaccine did not show decreased ability to neutralise a recombinant virus with 3 key B.1.1.7 S mutations; Δ69-70 + N501Y + D614G (Xie et al., 2021). Against authentic SARS-CoV-2, convalescent and BNT162b2 vaccine serum titres were equivalent against a mouse-adapted strain with N501Y and the parental strain (Rathnasinghe et al., 2021), whereas a full B.1.1.7 isolate showed a two-fold reduction in neutralisation by BNT162b2 vaccine sera (Diamond et al., 2021). Others have reported a 2-fold reduction in neutralisation of authentic B.1.1.7 virus by both convalescent and BNT162b2 vaccine sera with some convalescent sera which weakly neutralise the WT virus falling below the threshold of detection against B.1.1.7 (Skelly et al., 2021). Taken together, the evidence to date shows that the B.1.1.7 variant is equivalently or slightly less well neutralised by polyclonal sera and does not yet present a substantial risk of escape from pre-existing or vaccine-induced antibody immunity.

However, other than investigating its antigenicity, no studies have yet reported on the virological characteristics of the B.1.1.7 variant which might contribute to its emergence and spread. In this study, we experimentally characterised a panel of B.1.1.7 isolates. We tested whether B.1.1.7 shows enhanced replication compared to contemporaneous strains in Vero cells or in primary human airway epithelial cells grown at air liquid interface. We investigated whether B.1.1.7 lineage viruses have differences in their furin cleavage efficiency. Finally, we also tested whether different isolates of B.1.1.7 escape neutralisation by sera from convalescent and vaccinated patients. Collectively our data suggest that neither immune escape nor increased replication capacity account for the rapid emergence and increased transmission of B.1.1.7, and instead point to an increase in furin cleavage of spike that may enhance infectiousness of the variant.

## Results

### SARS-CoV-2 lineage B.1.1.7 replicate poorly and displays a small plaque phenotype in Vero cells

To characterise SARS-CoV-2 lineage B.1.1.7 (Variant of Concern 202012/01), we assembled a panel of isolates including five B.1.1.7 lineage viruses derived from four independent patients, together with historic and contemporaneous isolates from other SARS-CoV-2 lineages. Full sequence names and lineage information are provided in Methods. For each isolate, we verified the sequence and established the genome content and infectivity of the Vero cell passage 2 stock. During viral titration, we noticed that the B.1.1.7 isolates displayed a small plaque phenotype in Vero cells compared to non-B.1.1.7 viruses (Figure 1A). The mode area of plaques produced by the historic WT SARS-CoV-2 isolate (IC19) collected from a patient in March 2020 and containing D614G, was ≥ 0.10 pixel^2^(x10^3^), whereas the B.1.1.7 variants produced plaques of ≤ 0.06-0.08 pixel^2^(x10^3^) (Figure 1B, top panel). Plaques of contemporaneous non-B.1.1.7 isolates; B.1.258 with Δ69-70 and N439K spike mutations, and B.1.117.19 carrying an A222V mutation (Figure 1B, lower panel) did not significantly differ from WT IC19.

**Figure 1.**
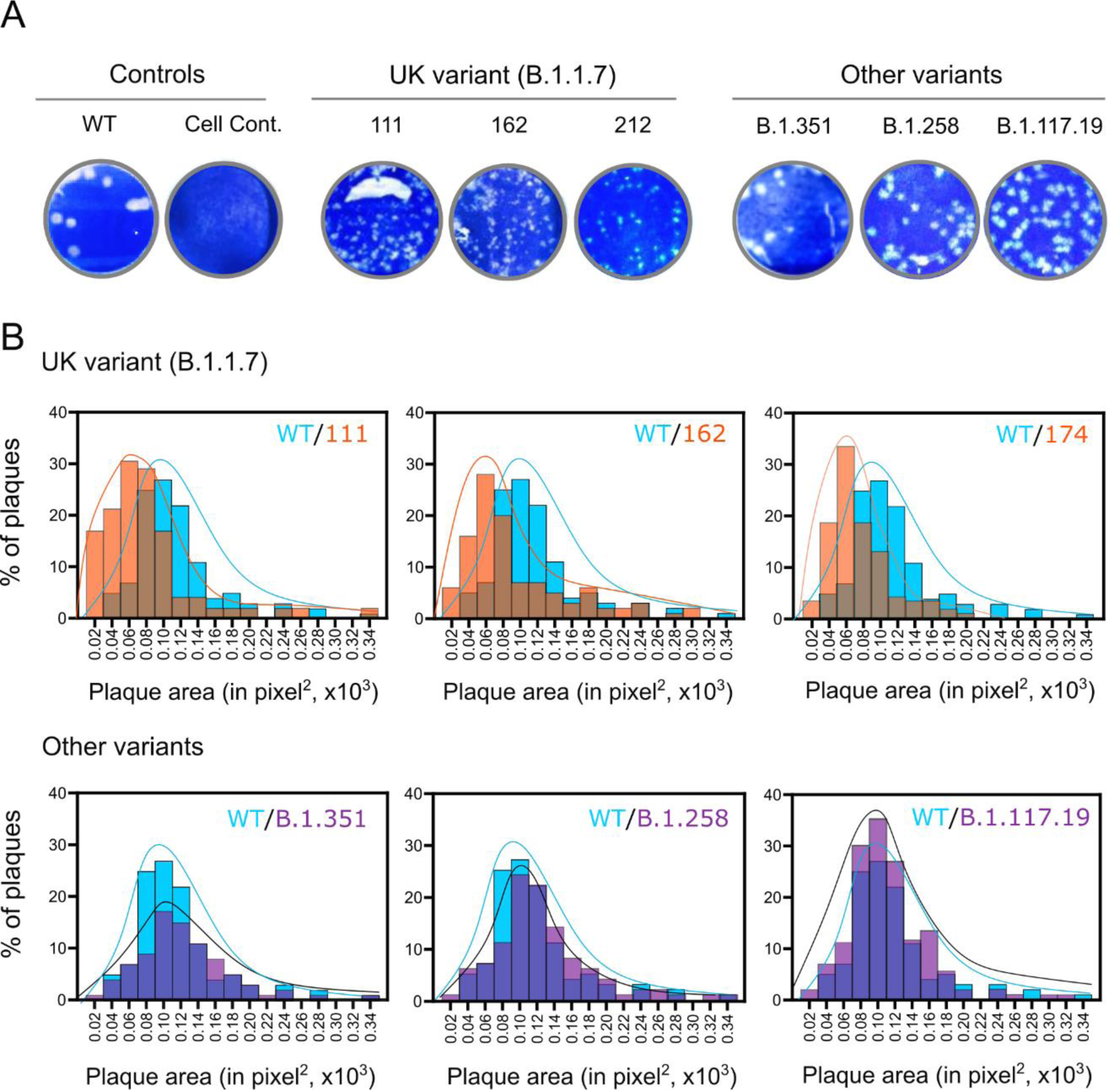
Plaque sizes of SARS-CoV-2 lineage B.1.1.7 isolates. Vero cells were infected with WT SARS-CoV-2 (IC19) or the indicated B.1.1.7 or other contemporaneous variants. Cells were overlaid with agar at 37°C for 72 h and plaques were visualised through crystal violet staining. (A) Representative images of virus plaques. (B) Histograms of plaque sizes quantified using ImageJ. A minimum of 300 plaques per virus were measured.

### Increasing prevalence of SARS-CoV-2 lineage B.1.1.7 is not accounted for by a replication advantage over non-B.1.1.7 viruses in a human airway epithelial cell model

To investigate whether increased transmissibility of the B.1.1.7 variant could be explained by more rapid replication kinetics, we infected primary human airway epithelial (HAE) cells or Vero cells at a standardised multiplicity of infection. In a first experiment, we normalised the virus inputs based on infectivity as measured by plaque assay on Vero cells. We infected Vero or HAE cells at 0.01 plaque forming units (pfu)/cell and quantified viral replication by assaying virus in Vero cell media or in extracellular washes obtained from the apical surface of infected HAE cells collected at different time points after infection, by qPCR for E gene (Supplementary 1). Levels of B.1.1.7 genomes were higher at all timepoints relative to the pair of non-B.1.1.7 viruses tested (B.1.117.19 and WT IC19). The increase genome level at time 0 is consistent with the high genome:pfu ratio in the B.1.1.7 stock. The increased genome copies of B.1.1.7 at later timepoints in HAE cells might also be accounted for by the higher input since titration on Vero cells had underestimated the infectivity of the stock. In contrast, in Vero cells, despite its higher input levels at time 0, B.1.1.7 showed a growth defect relative to B.1.117.19 and IC19 isolates.

As calculating viral titre on Vero cells had underestimated B.1.1.7 infectivity, we next normalised inputs based on genome copies as measured by E gene qPCR. HAE or Vero cells were infected with 1×10^4^ genomes/cell of B.1.1.7 alongside contemporaneous B.1.258 and B.1.117.19 isolates and again virus released from infected cells was quantified by qPCR (Figure 2A). In Vero cells, the B.1.1.7 isolate again displayed a significant growth defect. However, in HAE cells, the three isolates showed no significant difference in growth kinetics. To further confirm the differential growth phenotype of B.1.1.7 in different cell types was consistent, we infected Vero or HAE cells with a different B.1.1.7 isolate alongside the B.1.258 strain (Figure 2B). Again, a significant defect was seen for B.1.1.7 replication in Vero cells but there was no difference in replication kinetics in HAE cells. Transmission electron microscopy of fixed HAE sections at 72 hours post-infection showed no obvious differences in the morphology of B.1.1.7 virions compared to those of the other isolates (Figure 2C).

**Figure 2.**
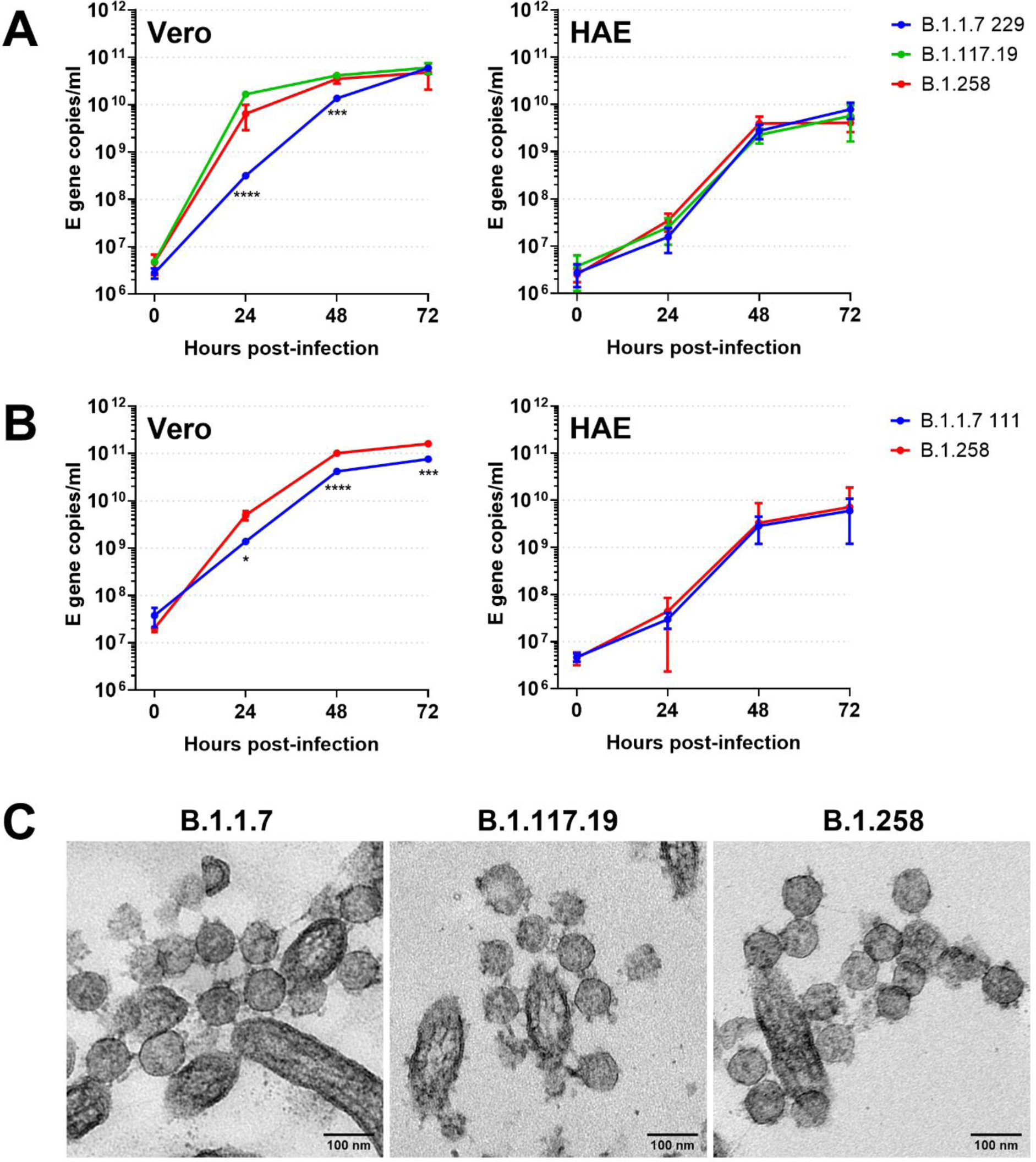
Comparative replication kinetics of SARS-CoV-2 lineage B.1.1.7 isolates in Vero and primary human airway epithelial (HAE) cells. Triplicate wells of Vero or primary human airway epithelial cells were infected with SARS-CoV-2 isolates at a multiplicity of 1×10^4^ genomes/cell and replicating virus released in media or washed from HAE apical surface was quantified by E gene qPCR at time points post-infection. (A) B.1.1.7 isolate 229 replication in Vero cells relative to non-B.1.1.7 B.1.117.19 and B.1.258 isolates (left hand panel) and in HAE cells (right hand panel). (B) B.1.1.7 isolate 111 and B.1.258 isolate replication in Vero cells (left hand panel) and in HAE cells (right hand panel). Statistical differences measured by ANOVA on log transformed data. *, P<0.05; ***, P<0.001; ****, P<0.0001. (C) TEM images of extracellular virions at the HAE apical cell surface at 72 hours post-infection.

### The P681H substitution in the B.1.1.7 spike confers an optimised S1/S2 furin cleavage site

We next investigated whether the phenotype of attenuated replication in Vero cells we observed for B.1.1.7 could be explained by differences in the efficiency of cleavage of its spike (S) surface protein since one of the lineage defining mutations, P681H, is close to the furin cleavage site. We and others have previously shown that efficiency of S1/S2 cleavage in producer cells can modulate the entry efficiency of SARS-CoV-2 into different cell types (Hoffmann, Kleine-Weber, & Pöhlmann, 2020; Johnson et al., 2021; Peacock et al., 2020). Deleting the furin cleavage site enhances entry into cell lines that lack TMPRSS2 protease expression (e.g. Vero) but attenuates entry into TMPRSS2-expressing cell lines (e.g. HAE or Calu-3 cells). We hypothesised that the loss of replication we observed in Vero cells might be accounted for by increased furin cleavage of the B.1.1.7 S, resulting in further virus instability. To test this, we generated a lentiviral pseudotype (PV) bearing full B.1.1.7 S, or S with P681H alone, and assessed the efficiency of cleavage by western blot (Figure 3A). The B.1.1.7 S showed increased cleavage compared to WT D614G S, more akin to that of D614G S containing a highly optimized polybasic furin cleavage site from an H5N1 avian influenza haemagglutinin (H5CS), as previously described (Peacock et al., 2020). Of the 9 mutations present in the B.1.1.7 S, P681H alone was sufficient to confer the optimised cleavage in line with its proximity to the S1/S2 cleavage site. To confirm the cleavage phenotype for authentic SARS-CoV-2 virus, we assessed S cleavage efficiency of B.1.1.7 isolate 229 and WT D614G (IC32) virus by western blot (Figure 3B). Again, increased cleavage was observed for B.1.1.7 S compared to WT D614G S.

**Figure 3.**
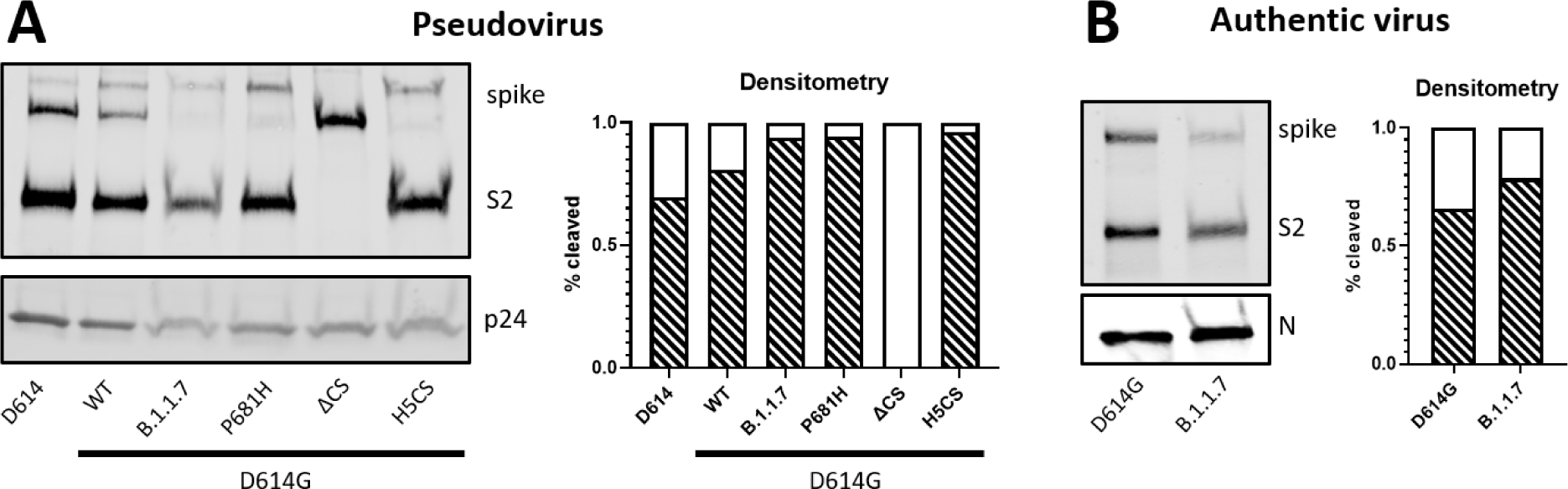
SARS-CoV-2 lineage B.1.1.7 spike cleavage. (A) Lentiviral pseudotypes bearing various SARS-CoV-2 spike (S) glycoproteins with the indicated mutations were concentrated by ultracentrifugation and the proportion of cleaved S quantified by densitometry on western blot. ΔCS - furin cleavage site removed; H5CS – influenza H5 haemagglutinin polybasic cleavage site. (B) S cleavage of concentrated, authentic SARS-CoV-2 D614G and B.1.1.7 lineage viruses and quantification by densitometry.

### SARS-CoV-2 lineage B.1.1.7 is susceptible to human convalescent sera from individuals infected in the first pandemic wave

The potential escape of newly arising SARS-CoV-2 variants from neutralising antibodies raised against previous strains is a global concern, particularly with vaccine efforts accelerating in many regions. To test whether B.1.1.7 lineage escapes from neutralising antibodies, we carried out live virus neutralisation assays to assess sera neutralisation of B.1.1.7 isolate 229 with WT IC19 as a comparator. Neutralisation was carried out with sera collected from healthcare workers (HCW) in May 2020 collected at least 21 days since a mild or asymptomatic SARS-CoV-2 infection confirmed by PCR (n=8), and sera from individuals in December 2020/January 2021 with previous sequence-confirmed B.1.1.7 infection (n=8) (Figure 4A). All sera were tested against approximately 100 TCID_50_ of the two virus isolates in the same assay and NT_50_ values were calculated. Virus inputs for the assay were approximately equivalent based on the back titration carried out concurrent with the neutralisation assay (Figure 4A). The median titres of the May 2020 sera against B.1.1.7 and IC19 were 87 and 63 respectively, with 6/8 (75%) showing a less than 2-fold difference in their ability to neutralise the two viruses. The remaining two May 2020 sera showed 2.0-fold and 2.6-fold increased ability to neutralise B.1.1.7. Overall, no significant reduction in titre of May 2020 sera was observed against the B.1.1.7 isolate indicating a lack of antibody escape. Sera collected following infection with B.1.1.7 had a median titre of 132 against the B.1.1.7 isolate and 35 against IC19. All 8 (100%) of these sera showed a greater than 2-fold (2.0-fold to 5.7-fold) increased ability to neutralise the homologous B.1.1.7 virus over IC19. This suggests that antibody responses mounted against SARS-CoV-2 lineage B.1.1.7 viruses retain efficacy against previously circulating strains but that this response is reduced perhaps owing to an immunodominant response to B.1.1.7-specific epitopes. To complement the findings of the authentic virus neutralisation assay, lentiviral PV bearing the S glycoprotein of either WT or B.1.1.7 virus were tested with a subset of the May 2020 and B.1.1.7 sera (Figure 2B). The same trend was observed with May 2020 antisera showing no significant difference in neutralisation, but B.1.1.7 convalescent antisera more efficiently neutralising B.1.1.7 spike-bearing PV.

**Figure 4.**
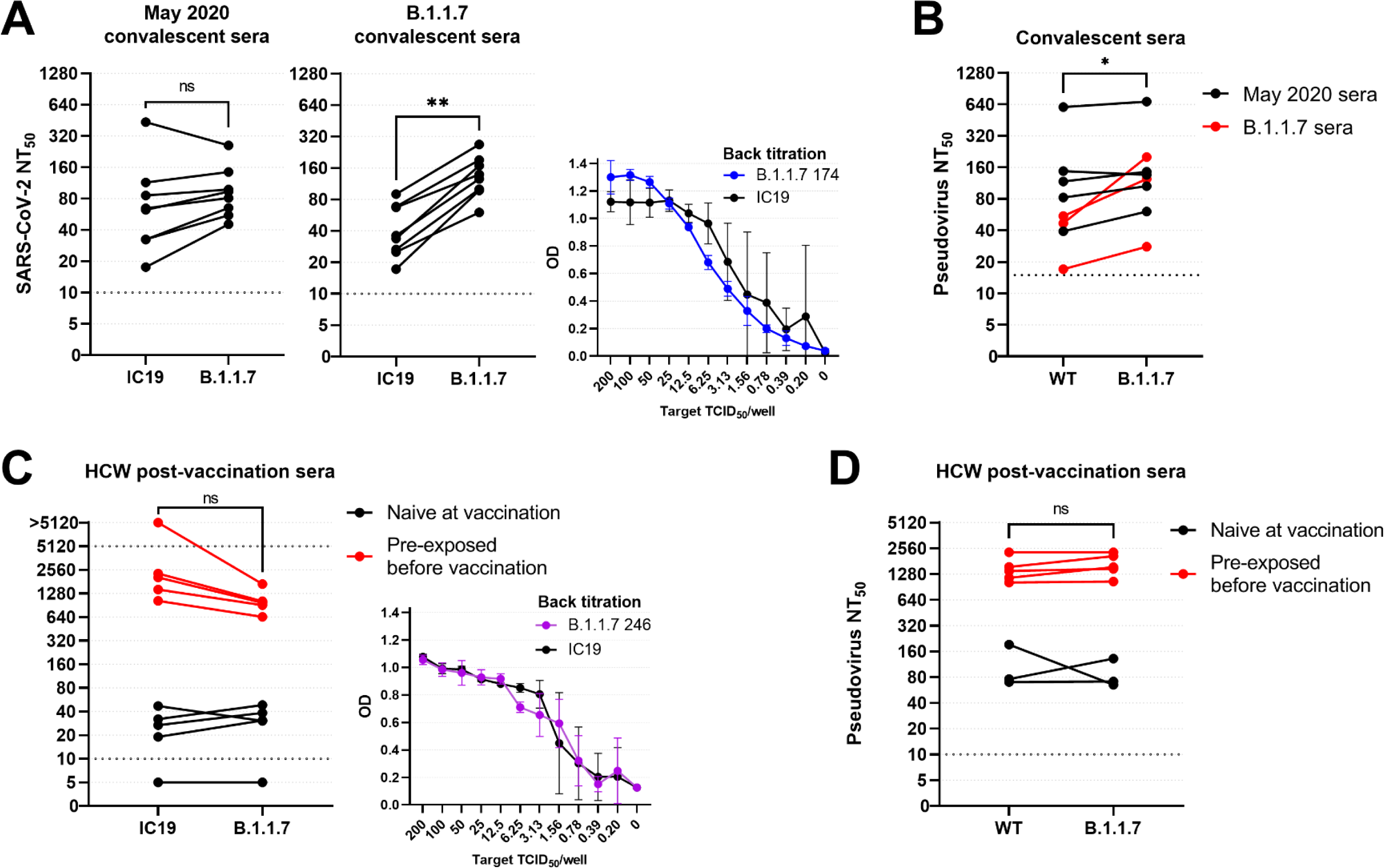
Neutralisation of live SARS-CoV-2 lineage B.1.1.7 virus by convalescent and post-vaccination sera. (A) Sera raised against early pandemic viruses in May 2020 (n=8) and B.1.1.7 lineage viruses (n=8) were assessed for their ability to neutralise B.1.1.7 isolate 229 and WT IC19 control virus by live neutralisation assay. NT_50_ values were calculated for each serum sample against both viruses. There was no significant difference in the ability of early pandemic sera to neutralise the two viruses but B.1.1.7 sera were significantly better able to neutralise the homologous lineage virus than IC19. Wilcoxon matched-pairs signed rank test was used for comparisons. A 2-fold dilution series of the virus input used for the neutralisation assay was performed and the mean OD of 4 replicate wells is shown indicating that approximately equivalent amounts of each virus were used as challenge. (B) NT_50_ values for May 2020 sera (n=5) and B.1.1.7 sera (n=3) against lentiviral pseudotypes bearing WT or B.1.1.7 spike. (C) Sera taken from healthcare workers (HCW) 21-25 days after receiving their first BNT162b2 (Pfizer-BioNTech) vaccine dose were assessed for their ability to neutralise B.1.1.7 isolate 246, relative to IC19. HCW sera were divided into those who had been exposed to SARS-CoV-2 prior to receiving the vaccine (red lines, n=5) and those who were naïve when receiving the vaccine (black lines, n=5), as determined by Fortress lateral flow immunoassay test. (D) NT_50_ values for pre-exposed vaccine sera (n=5) and naïve vaccine sera (n=3) against WT and B.1.1.7 pseudotypes.

### BNT162b2 vaccination induces equivalent neutralising antibody responses against B.1.1.7

To investigate whether antibody responses elicited by currently distributed vaccines were also likely to be protective against B.1.1.7, sera were taken from HCW on the day of receiving their first dose of the Pfizer/BioNTech BNT162b2 vaccine (day 0) and approximately 21-25 days post-vaccination. Day 0 sera were tested by Fortress lateral flow immunoassay (LFIA) for the presence of SARS-CoV-2-specific antibodies to segregate HCW into individuals who were pre-exposed to virus (n=5) and those who were naïve on the day of vaccination (n=5). The post-vaccination sera were then assessed for neutralising antibodies against B.1.1.7 isolate 246 and IC19 (Figure 4C). Neutralising antibody titres post-vaccination were significantly higher in sera from pre-exposed (red lines) compared to naive HCW (black lines) with median titres against IC19 of 2038 and 32 respectively. Post-vaccination sera from pre-exposed HCW also neutralised B.1.1.7 well with a median titre of 968 although 3/5 sera showed a greater than 2-fold decrease (2.1-fold to 3.0-fold) in their ability to neutralise B.1.1.7 compared to IC19. This drop in titre against B.1.1.7 was not observed in the PV neutralisation assay with median titres of 1392 against IC19 and 1540 against B.1.1.7 (Figure 4D). For HCW who were naïve on the day of vaccination, titres were low post-vaccination and there was no significant difference in their responses to B.1.1.7 and IC19 (Figure 4C, D). The four sera which were above the threshold of detection of the live virus neutralisation assay against both B.1.1.7 and IC19 had median titres of 34 and 29 respectively.

## Discussion

SARS-CoV-2 lineage B.1.1.7 (Variant of Concern 202012/01) poses a clear threat to ongoing efforts to control the burden of the COVID-19 pandemic due to its increased transmissibility and association with an increased hazard of death. B.1.1.7 has rapidly risen to predominate in the UK and beyond but the virological traits which have allowed it to do this have yet to be described. Here, we show that B.1.1.7 does not demonstrate increased replication in human airway epithelial (HAE) cells. This suggests that increased viral replication is not responsible for the increase of viral transmission seen in the B.1.1.7 lineage. This differs from viruses with the D614G mutation in spike (S), whose global spread was at least partially accounted for by increased viral replication that was evident in HAE cultures (Hou et al., 2021). In infected individuals, a small but significant decrease in Ct values in G614 compared to D614 infections in the UK demonstrated increased viral load for variants carrying D614G (Volz, Hill, et al., 2021). For B.1.1.7, early analysis using the number of mapped sequencing reads or Ct values as proxies for viral load suggested an increased viral burden associated with B.1.1.7 infection (Golubchik et al., 2021; Kidd et al., 2020) but more recent reports show no such association at either the population level or longitudinally in individuals (Kissler et al., 2021; Walker et al., 2021). The more recent reports and our data suggest there is no clear replicative advantage for B.1.1.7 *in vitro* or at the population level.

In contrast to the equivalent replication we saw in HAE cells, B.1.1.7 showed reduced replication and a corresponding smaller plaque size on Vero cells compared to other tested viral lineages (Figures 1 and 2). Our previous work highlighted that S1/S2 cleavage in the producer cell, conferred by a polybasic stretch at the cleavage site, is advantageous in cells expressing abundant TMPRSS2 but deleterious in cells lacking TMPRSS2 such as Vero cells (Peacock et al., 2020). Analysis here revealed that the P681H substitution directly adjacent to the S1/S2 cleavage site resulted in increased cleavage of S (Figure 3) and could be responsible for the reduced entry into Vero cells. However, increased cleavage as an explanation for the deleterious effect in Vero cells was not accompanied by a corresponding advantage in HAE cells. It is possible that increased cleavage could be beneficial to transmission and entry into the human airway, but that this phenotype is not observed when infecting HAE cultures with a relatively large amount of virus compared to the likely dose during transmission. Further experiments to establish the lowest dose required to initiate infection in vitro and in vivo may clarify this. Additionally, the decreased growth on Vero cells means that care must be taken when growing B.1.1.7 viral stocks since deletion or mutation of the S1/S2 cleavage site might be selected for in these cells. Viral stocks should be sequenced to confirm that the cleavage site remains intact. In addition, viral titres calculated on Vero cells may lead to an underestimation of the number of infectious viruses due to the high pfu:genome ratio.

We confirmed that B.1.1.7 does not escape from antibody immunity after natural infection with previously circulating variants, or vaccine expressing the S protein of older variants, suggesting that immune escape does not account for increased transmission (Figure 4). This is unsurprising given that there is no observed increase in reinfection reported with B.1.1.7 nor was there high seroprevalence in the UK during the emergence of this variant. Our finding is supported by many studies using B.1.1.7 viruses and PV which show either no reduction or a modest reduction in polyclonal serum titres (Collier et al., 2021; Diamond et al., 2021; Hu et al., 2021; Muik et al., 2021; Rees-Spear C et al., 2021; Wang et al., 2021; Wu et al., 2021). There is some evidence of heterogeneity of responses and neutralising titres for some individuals with initially low responses against WT virus can drop below limits of detection against the B.1.1.7 variant (Skelly et al., 2021). However, since the correlate of protection is not yet established for SARS-CoV-2, the significance of this drop is still not clear. Whilst the N501Y mutation has been implicated in loss of binding of some RBD targeting monoclonal antibodies, and B.1.1.7 also contains mutations in the N-terminal domain (NTD) of S which may allow for escape against NTD targeting antibodies, the small effect on neutralisation titres by polyclonal sera is reassuring and suggests further antigenic domains on S that contribute to protection. In our study, the reduced ability of sera raised against B.1.1.7 to neutralise the historic IC19 virus relative to the homologous virus (Figure 4A) may suggest that immune responses to B.1.1.7 are more focussed meaning B.1.1.7 S may not be a preferred choice as the basis of future vaccine updates.

B.1.1.7 is one of a growing number of variants of concern which are showing increased transmission. These variants share many convergent or parallel mutations of S as well as in NSP6 and ORF8 suggesting that these mutations are likely adaptive and likely multiple mutations are needed for an increase in transmissibility. It is notable that several emerging variants contain mutations which could increase cleavage of S. These include other mutations adjacent to the cleavage site such as P681R, Q677H, and Q677P (Hodcroft et al., 2021) as well as further mutations (H655Y, A701V) which are more distant in primary sequence but proximal to the furin cleavage site in the 3D structure of S. We therefore hypothesise that an increase in the efficiency of furin cleavage is an important contributing factor to the increase in transmission of these variants. The B.1.1.7 lineage continues to evolve and it is notable that several isolates have gained additional S mutations such as E484K which has been shown to cause a 9.6-fold decreased neutralisation by vaccine sera in a B.1.1.7 background (Collier et al., 2021), and which could further increase receptor binding avidity in combination with N501Y. E484K and N501Y together have been associated with other rapidly emerging variants of the B.1.351 and P.1 lineages.

It is likely that the increased transmissibility of B.1.1.7 owes to subtle optimisation and balancing of a number of virological traits which include and extend beyond those investigated here. Once the genetic determinants of increased transmissibility of the B.1.1.7 variant are found it will be important to remain vigilant for these hallmarks in other emerging variants, especially if in combination with mutations which confer vaccine escape.

## Materials and Methods

### Cells

African green monkey kidney (Vero) cells (Nuvonis Technologies) were maintained in OptiPRO SFM (Life Technologies) containing 2X GlutaMAX (Gibco). Human embryonic kidney cells (293T) were maintained in Dulbecco’s modified Eagle’s medium (DMEM), 10% fetal calf serum (FCS), 1% non-essential amino acids (NEAA), 1% penicillin-streptomycin (P/S). All primary and continuous cell lines were maintained at 37°C, 5% CO_2_. 293T-ACE2 cells were generated as previously described (Peacock et al., 2020; Rebendenne et al., 2021) and were maintained with 293T media supplemented with 1 μg/ml of puromycin. Primary nasal human airway epithelial (HAE) cells at air-liquid interface (ALI) were purchased from Epithelix for the experiment in Figure 1B and primary bronchial HAE cells from Epithelix for the experiment in Supplementary 1. The basal MucilAir medium (Epithelix) was changed every 2-3 days for maintenance of HAE cells. For experiment 1A, HAE cells at ALI were differentiated in-house as follows under Health Research Authority study approval (REC ref: 20/SC/0208; IRAS: 282739). A nasal brushing of the turbinate was acquired using 3-mm bronchial cytology brush and placing the biopsy into warm PneumaCult-Ex Plus Medium (STEMCELL Technologies, Cambridge, UK). The cells were dissociated from the brush by gentle agitation before seeding into a single well of a collagen (PureCol from Sigma Aldrich) coated plate. Once confluent the cells were passaged and expanded further in a flask before passaging a second time and seeding onto transwell inserts (6.5 mm diameter, 0.4 μm pore size, Corning) at a density of 24,000 cells per insert. Cells were cultured in PneumaCult-Ex Plus (STEMCELL Technologies, Cambridge, UK) until confluent, at which point the media was replaced with PneumaCult-ALI in the basal chamber and apical surface exposed to provide an air liquid interface (ALI). Cilia were observed between 4-6 weeks post transition to ALI.

### Serum samples

Sera were collected under ethical approval as stated in the Ethical Approval and heat-inactivated at 56°C for 30 minutes before use in assays.

### Viruses

SARS-CoV-2 infectious swabs were collected as approved in Ethical Approval. Viruses were isolated by inoculating 100ul of neat swab material onto 24-well plates of Vero cells, incubating at 37°C, 5% CO_2_ for 1 hour before adding 1 ml OptiPRO SFM supplemented with 2X Glutamax, 1% P/S and 1% amphotericin and incubating again for 5-7 days until cytopathic effect was observed. Isolates were passaged twice in Vero cells and used for subsequent experiments. For western blot analysis, virus supernatants were concentrated by spinning through an Amicon^®^ Ultra-15 Centrifugal Filter Unit followed by an Amicon^®^ Ultra-0.5 Centrifugal Filter Unit with 50 kDa exclusion size.

| Name used in text | Virus name | GISAID Accession ID | PANGO lineage |
| --- | --- | --- | --- |
| **B.1.1.7 111** | hCoV-19/England/205090256/2020 | EPI_ISL_747517 | B.1.1.7 |
| **B.1.1.7 162** | hCoV-19/England/205080329/2020 | EPI_ISL_722999 | B.1.1.7 |
| **B.1.1.7 212** | hCoV-19/England/204690005/2020 | EPI_ISL_693401 | B.1.1.7 |
| **B.1.1.7 229** | hCoV-19/England/204661721/2020 | EPI_ISL_693400 | B.1.1.7 |
| **B.1.1.7 246** | hCoV-19/England/205080610/2020 | EPI_ISL_723001 | B.1.1.7 |
| **B.1.117.19** | hCoV-19/England/204501194/2020 | EPI_ISL_660788 | B.1.117.19 |
| **B.1.258** | hCoV-19/England/204501206/2020 | EPI_ISL_660791 | B.1.258 |
| **B.1.351** | hCoV-19/England/205280030/2020 | EPI_ISL_770441 | B.1.351 |
| **IC19** | hCoV-19/England/IC19/2020 | EPI_ISL_475572 | B.1.13 |
| **IC32** | hCoV-19/England/IC32/2020 | n/a | B.1.238 |

### Determination of plaque size

Virus inoculum were serially diluted in serum-free DMEM (1:10) supplemented with 1% NEAA and 1% P/S, and added to Vero cell monolayers at 37°C for 1 h. Inoculum was then removed and plates were overlaid with DMEM containing 0.2% w/v bovine serum albumin, 0.16% w/v NaHCO_3_, 10 mM HEPES, 2mM L-Glutamine, 1x P/S, and 0.6% w/v agarose at 37°C for 3 d. The overlay was then removed, and monolayers were stained with crystal violet solution. Plates were then washed with tap water, dried and scanned on a flatbed office scanner at 600 dots per inch (dpi). Images were analysed using the Analyze Particles module of Fiji (ImageJ). Areas of ≥ 300 plaques were measured for each virus variant and expressed in pixels^2^. Virus plaque sizes were compared using the Kolmogorov– Smirnov test with Bonferroni correction (Prism 9.0; GraphPad). *P* ≤0.05 significant difference from WT.

### Virus growth kinetics

All dilution of viruses, wash steps and harvests were carried out with OptiPRO SFM (Life Technologies) containing 2X GlutaMAX (Gibco). For HAE cells, all wash and harvest steps were performed by addition of 200ul SFM and incubation for 10 mins at 37°C before removing SFM. To infect, basal medium was replaced, cells were washed once with SFM to remove mucus before addition of inoculum and incubation for 1 h at 37°C. Inoculum was removed, cell washed twice and the second wash taken as harvest for 0 hpi. For infection of Vero cells, overnight growth medium was removed, inoculum added and incubated for 1 h at 37°C before removal, two washes and replacement with 3ml SFM from which harvests were taken at timepoints.

### Conventional transmission electron microscopy (TEM)

HAE cells were fixed by placing them in 2.5% glutaraldehyde in 0.05M sodium cacodylate buffer at a pH 7.4 and left for 2 days at room temperature. Subsequently, the samples were incubated in 1% aqueous osmium tetroxide for 1 h at RT before en bloc staining by placing them in undiluted UA-Zero (Agar Scientific) for 30 minutes at RT. The samples were dehydrated using increasing concentrations of ethanol (50%, 70%, 90%, 100%), followed by propylene oxide and a mixture of propylene oxide and araldite resin (1:1). To embed the samples, they were placing in aradite and left at 60°C for 48 h. Ultrathin sections were cut using a Reichert Ultracut E ultramicrotome and stained using Reynold’s lead citrate for 10 minutes at RT. Images were acquired on a JEOL 1400Plus transmission electron microscope fitted with an Advanced Microscopy Technologies (AMT) XR16 charge coupled device (CCD) camera.

### E gene qPCR

RNA was extracted from virus supernatants using QIAsymphony DSP Virus/Pathogen Mini Kit on the QIAsymphony instrument (Qiagen). qPCR was then performed using AgPath RT-PCR (Life Technologies) kit on a QuantStudio(TM) 7 Flex System with the primers for E gene used in (Corman et al., 2020). A standard curve was also generated using dilutions viral RNA of known copy number to allow quantification of E gene copies in the samples from Ct values. E gene copies per ml of original virus supernatant were then calculated.

### Live virus neutralisation assay

The ability of sera to neutralise SARS-CoV-2 virus was assessed by neutralisation assay on Vero cells. Sera were serially diluted in OptiPRO SFM (Life Technologies) and incubated for 1 h at RT with 100 TCID50/well of SARS-CoV-2/England/IC19/2020 and transferred to 96-well plates pre-seeded with Vero-E6 cells. Serum dilutions were performed in duplicate. Plates were incubated at 37°C, 5% CO_2_ for 42 h before fixing cells in 4% PFA. Cells were treated with methanol 0.6% H2O2 and stained for 1 h with a 1:3000 dilution of 40143-R019 rabbit mAb to SARS-CoV-2 nucleocapsid protein (Sino Biological). A 1:3000 dilution of sheep anti-rabbit HRP conjugate (Sigma) was then added for 1 h. TMB substrate (Europa Bioproducts) was added and developed for 20 mins before stopping the reaction with 1M HCl. Plates were read at 450nm and 620nm and the concentration of serum needed to reduce virus signal by 50% was calculated to give NT50 values.

### Pseudovirus assays

SARS-CoV-2 spike-bearing lentiviral pseudotypes (PV) were generated as previously described (Peacock et al., 2020). PV for western blot analysis were concentrated by ultracentrifugation at 100,000 x g for 2 hours over a 20% (w/v) sucrose cushion. PV neutralisation assays were performed by serially diluting sera in 293T growth media and incubating for 1 h at 37°C with equal concentrations of PV. The PV/antisera mix was then added onto 293T-ACE2 cells. Serum dilutions were performed in duplicate. 293T-ACE2 were transduced for 48 hours before lysis with reporter lysis buffer (Promega). Luciferase luminescence was read on a FLUOstar Omega plate reader (BMF Labtech) using the Luciferase Assay System (Promega).

### Western blot analysis

Concentrated PV or virus was mixed with 4x Laemmli sample buffer (Bio-Rad) with 10% β-mercaptoethanol and run on SDS-PAGE gels. After semi-dry transfer onto nitrocellulose membrane, membranes were probed with mouse anti-p24 (abcam; ab9071), rabbit anti-SARS spike protein (NOVUS; NB100-56578) or rabbit anti-SARS-CoV-2 nucleocapsid (SinoBiological; 40143-R019). Near infra-red (NIR) secondary antibodies, IRDye^®^ 680RD Goat anti-mouse (abcam; ab216776) and IRDye^®^ 800CW Goat anti-rabbit (abcam; ab216773) were subsequently used to probe membranes. Western blots were visualised using an Odyssey Imaging System (LI-COR Biosciences).

## Ethical approval

Convalescent sera from healthcare workers at St. Mary’s Hospital at least 21 days since PCR-confirmed SARS-CoV-2 infection were collected in May 2020 as part of the REACT2 study with ethical approval from South Central Berkshire B Research Ethics Committee (REC ref: 20/SC/0206; IRAS 283805). Patient swabs for virus isolation and sera raised against B.1.1.7 were collected by the PHE Virology Consortium. The investigation protocol was reviewed and approved by the PHE Research Ethics and Governance Group and Incident Management team. PHE has legal permission, provided by Regulation 3 of the Health Service (Control of Patient Information) Regulation 2002, to process patient confidential information for national surveillance of communicable diseases. Further infectious swabs and sera were collected as part of the Assessment of Transmission and Contagiousness of COVID-19 in Contacts (ATACCC). Ethical approval for ATACCC was granted under the Integrated Network for Surveillance, Trials and Investigation of COVID-19 Transmission (INSTINCT; Ethics Ref: 20/NW/0231; IRAS Project ID: 282820) Sera collected from HCW 21-25 days after their first BNT162b2 vaccine dose were collected as part of a study approved by the Health Research Authority (REC ref: 20/WA/0123).

## Acknowledgements and Funding

Special thanks to members of the PHE Virology Consortium; Angie Lackenby, Shahjahan Miah, Steve Platt, Joanna Ellis, Maria Zambon and Christina Atchison, as well as PHE field staff for collection of infectious swabs, sequence information and serum samples used in this work. We thank the ATACCC Investigators including Ajit Lalvani and Jake Dunning (co-PIs), Robert Varro (Study Coordinator), Jessica Cutajaar (Senior Research Nurse), Joe Fenn, Rhia Kundu, Seran Hakki, Timesh Pillay (post-doctoral and clinician scientists), the ATACCC technicians, research nurses and support workers. We acknowledge the support of the NIHR Health Protection Research Unit in Respiratory Infections, Imperial College London (NIHR200927) and the ‘Assessment of Transmission and Contagiousness of COVID-19 in Contacts’ (ATACCC) grant from the Department of Health and Social Care (DHSC) COVID-19 Fighting Fund. Also thanks to Maria Prendecki and Michelle Willicombe for providing post-vaccination sera, Graham Cooke for REACT convalescent sera and Ranjit K. Rai and Paul Griffin for assistance with HAE cell culture. AS was supported by a British Society for Antimicrobial chemotherapy COVID-19 grant. This work was supported by the G2P-UK National Virology Consortium funded by UKRI.

---

**Supplementary 1.**
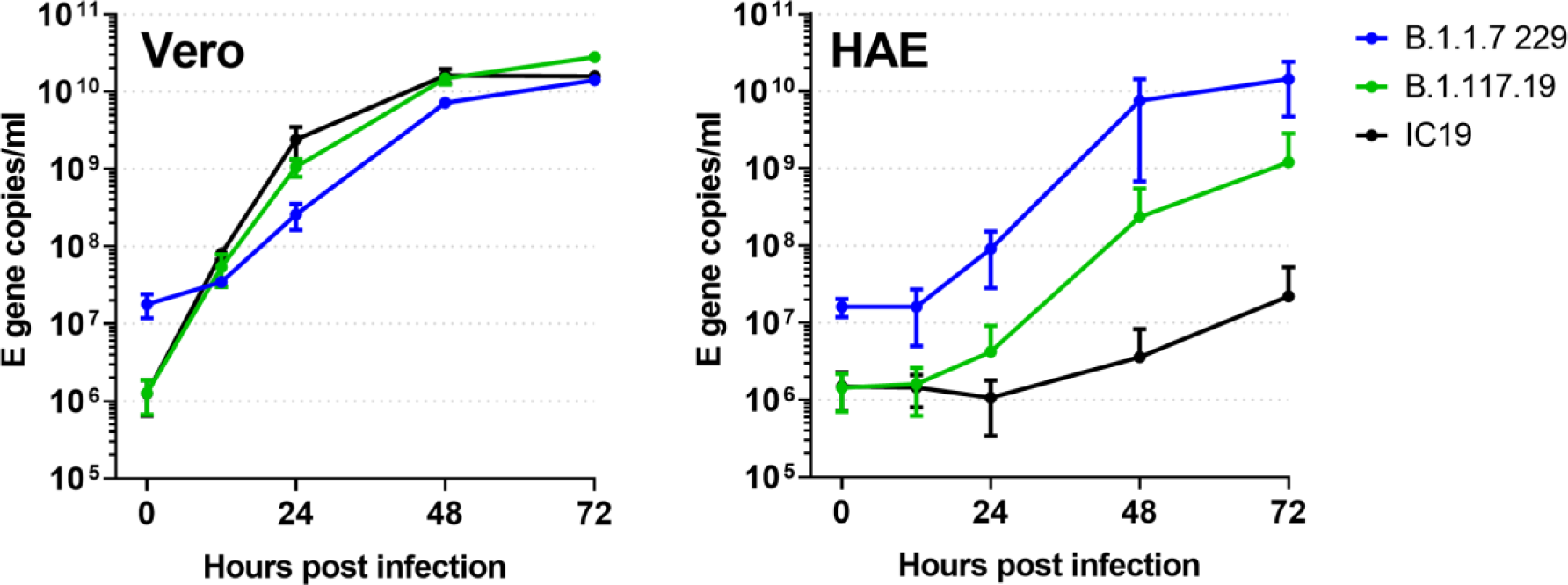
Replication kinetics of B.1.1.7 and non-B.1.1.7 isolates with input normalised by infectivity on Vero cells. Virus isolates were plaqued on Vero cells to determine infectious titre and then used to infect triplicate wells Vero or human airway epithelial (HAE) cells at a multiplicity of 0.01 pfu/cell. Replication was measured by E gene qPCR of Vero supernatants and HAE apical harvests. B.1.1.7 had higher genome input despite having an equivalent infectious dose as measured on Vero cells.

